# Roles of bacterial growth competition systems in colonization of the murine gut

**DOI:** 10.64898/2026.05.11.724384

**Authors:** Petra Muir, Jonas Kjellin, Evelina Kess, David Low, Sanna Koskiniemi

## Abstract

The gut microbiome is essential for human health. Although the gut microbiota is largely stable at the species level in healthy individuals, strain-level variation remains less understood. Many bacterial strains encode toxin delivery systems that may shape competition within the gut. Here, we investigate how contact-dependent growth inhibition (CDI) and colicins influence intestinal colonization by a competitive murine *Escherichia coli* isolate, R12. We show that R12 can colonize an intact mouse gut microbiota by displacing resident *Enterobacteriaceae*, but success depends on multiple interacting factors. CDI systems and colicins provide a competitive advantage against resident *E. coli*, particularly during early colonization, while metabolic flexibility and access to alternative carbon sources support long-term persistence. Colonization outcomes vary between hosts and are shaped by resident microbiota composition, strain-level competition, and the initial invader-to-resident ratio. Overall, successful gut invasion is determined by the combined effects of bacterial antagonistic systems, metabolic capacity, and ecological context.

## Main text

The gastrointestinal tract of mammals is colonized by a large number of microorganisms, commonly referred to as the normal gut microbiota. The composition of the gut microbiota for host extraction and synthesis of nutrients and metabolites, development of the immune system and preventing colonization of pathogens is increasingly recognized ^1^. Although the microbiome in healthy individuals remains relatively stable over time ^2^, changes on strain level are not well understood. Strain level variation in e.g. *E. coli* population are important risk factors for colon cancer (colibactin production) as well as urinary tract infections (uropathogenic strains) ^3-5^.

*De novo* colonization of the gut involves several challenges, including overcoming the colonization resistance provided by the resident microbiota ^6^. The mechanisms behind colonization resistance are not yet fully understood, but most likely involve nutrient competition ^7,8,9^ and toxin delivery from established bacteria ^10-17,18,19^. Bacterial toxins can be delivered either through secretion to the extracellular milieu, e.g. microcins and colicins, or through direct cell-to-cell toxin delivery as with contact-dependent growth inhibition (CDI) or type 6 secretion systems (T6SS), Reviewed in ^20,21,22^. Expression of specific immunity proteins protects toxin-expressing cells from inhibiting their own growth (self-recognition)^20,23,24^. Previous findings have shown that microcins and T6SS, with broad host-ranges, contribute to bacterial colonization ^11,19^. In contrast, although CDI was first identified in the gut commensal EC93, its role in gut colonization remains poorly understood ^25,26^.

CDI can be considered a short- and narrow-range weapon, as delivery of toxic effector proteins requires physical contact with targeted cells and interaction with specific outer-membrane proteins that determine host-range^27^. Also colicins have narrow host-range, using the same type of receptors as CDI, but as they are secreted to the extra-cellular milieu their range can be considered longer ^20^. To recognize the correct target, specific receptor binding domains (RBDs) in the CdiA and Colicin proteins are used for target cell recognition ^20,27-31^. Following engagement of the target cell receptors, the CdiA-CTs and colicin cytotoxic domains, are translocated into target cells ^32^ where they display a range of activities, including DNase, RNase and pore-forming functions^20,28,33-35^.

To improve our understanding of bacterial host colonialization, we studied the highly competitive *E.coli* isolate R12 from rat in a murine modell, identifying its toxin-delivery systems and their contribution to colonisation. We find that *E. coli* R12 is able to colonize an intact mouse gut microbiota, but that its success depends strongly on both its toxin delivery systems and the composition of the resident microbiota. Genome analysis shows that R12 encodes two CDI systems and three colicins, which are all functional and contribute to competitive fitness against other *E. coli* strains, particularly within the *Enterobacteriaceae* niche. *In vivo* experiments demonstrate that wild-type R12 outcompetes resident *E. coli* more effectively than mutants lacking CDI and/or colicins, especially during later stages of colonization. However, colonization success is highly variable between mice and is influenced by the initial invader-to-resident ratio, resident strain competition systems, and the presence of alternative metabolic capabilities. R12’s ability to utilize unique carbon sources and its metabolic flexibility likely support colonization independently of toxin systems, particularly at lower competitive advantages. In contrast, CDI and colicins are crucial for early dominance and successful invasion when competing with established *E. coli* strains. Overall, colonization is determined by an interplay of bacterial antagonistic systems, metabolic capacity, resident microbiota composition, and initial population ratios. Together, these findings highlight how metabolic fitness and contact-dependent antagonism synergize to drive colonization—offering a framework for designing targeted microbiome interventions and competitive bacterial therapeutics.

### *E. coli* R12 can colonize an occupied niche

To determine if *E. coli* R12 can overcome colonization resistance mediated by an intact normal intestinal microbiota, we inoculated female BALB/c mice with 10^7^ rifampicin (RIF)-resistant *E. coli* R12 by oral gavage and measured the prevalence of R12 in the gut by plating fecal material on RIF plates (Fig. 1A). *E. coli* R12 levels decreased for the first few days post infection (p.i.) to increase to around 10^6^ CFU on day 8 (Fig. 1B). This suggests that *E. coli* R12 is able to colonize the mouse gut without prior niche clearance. Previous studies have shown that the lack of *Enterobacteriaceae* in normal gut microbiota of mice greatly increases colonization efficacy of *Salmonella enterica* ^36^. However, all mice had *Enterobacteriaceae* in their feces (Fig. 1C), suggesting that *E. coli* R12 is a successful colonizer even when the intestinal niche is already occupied.

**Figure 1.**
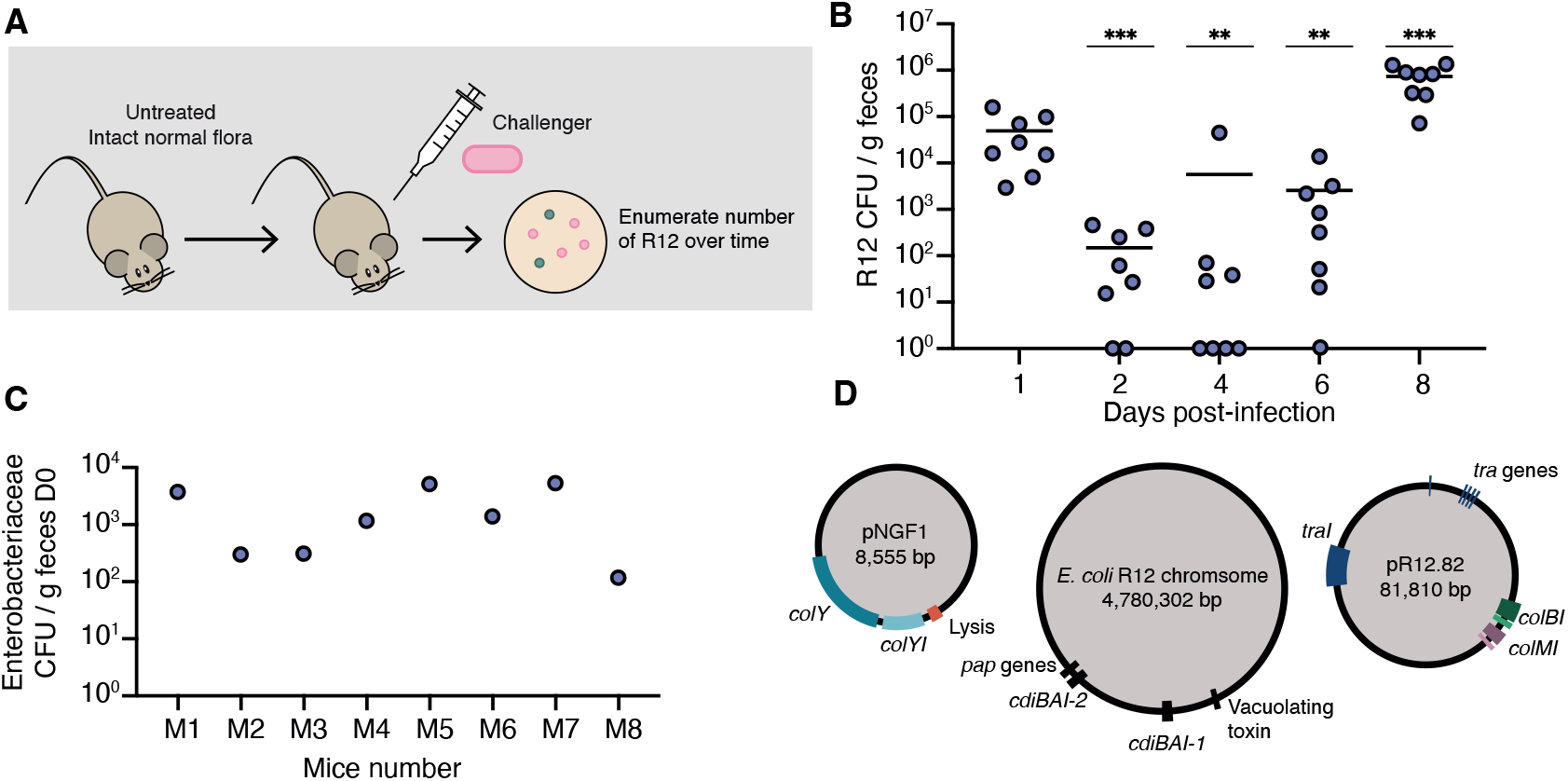
*E. coli* R12 colonization of mice. (A) Schematic overview of mouse experiment with intact gut microbiota. (B) Untreated BALB/c mice were inoculated orally with 10^7^ *E. coli* R12 and the number of CFU were determined by platin fecal material on MacConkey agar plates with or without Rifampicin. The data is plotted as CFU/g feces relative to day 1. Each dot represents one mouse (n = 8). P values were obtained using Mann-Whitney U test comparing the different days to day 1 p.i. *, p ≤ 0.05, ** p ≤ 0.01, *** p ≤ 0.001. (C) *Enterobacteriaceae* before the infection measured as cfu/ g feces on MacConkey agar plates . (D) Overview of the *E. coli* R12 genome. Genes encoding CDI systems (*cdiBAI*), colicins (*colBI, colMI, colYI*), transfer proteins (*tra*), P pili (*pap*) as well as a vacuolating toxin are indicated.

### Identification of the *E.coli* R12 toxin delivery arsenal

To investigate if toxin delivery systems could play a role in the ability of R12 to colonize an occupied niche, we performed whole genome sequencing (Pacific Biosciences). The sequences assembled into three contigs: a 4.8 Mb chromosome; a large 81.8 kb plasmid (pR12.82); and a small 8.6 kb plasmid (Fig. 1D) (accession: GCA_042192135.1). Bioinformatic analysis of the genome using Bagel 4^37^ revealed the presence of two chromosomal *cdiBAI* loci and three plasmid-encoded colicins, colicins B, M and Y (Fig. 1D).

The CdiA-proteins of the two identified CDI systems show high levels of homology (94.6% aa identity) differing only in their C-terminal toxin domains (Fig. S1). The RBDs of CdiA-1 and CdiA-2 are identical and highly homologous (99.3% identity) to the RBD of CdiA^UPEC536^ (Fig. S2), which recognizes OmpC/OmpF as receptor on target cells ^27,29^. Aligning the CdiA-CTs with toxins of known function suggests that *cdiA*^R12-CT1^ encodes an AcrB-dependent ionophore toxin as it is 100% identical to that of *E. coli* EC93 CdiA^EC93-CT2^ on protein level (Fig. S3) ^26^. The toxic activity of CdiA^R12-CT2^ is unknown as we were unable to find significant homology to previously described proteins.

The genes encoding colicin B and colicin M are located on an 82 kb plasmid (Fig. 1D). Colicins B and M are known to exploit siderophore receptors FepA and FhuA for target cell entry ^20^. Colicin B is a pore-forming colicin, causing depolarization of the cytoplasmic membrane, whereas colicin M inhibits peptidoglycan synthesis ^38-40^. R12 also contained the previously characterized 8.6 kb pColY plasmid. This plasmid was first isolated from the murine commensal *E. coli* MP1(accession: CP139039) ^41,42^ and later also found in NGF-1 (accession: CP016007))^43,44^, and encodes Colicin Y, another pore-forming colicin that recognizes OmpA as receptor ^45,46^. Bioinformatic analyses show that *E. coli* R12 is not MP1 (Table S1). Taken together, R12 encodes numerous bacterial toxin delivery systems that could be important for its colonization ability.

### *E. coli* R12 has two active CDI systems and produces two colicins

To determine whether CDI systems and colicins affect bacterial colonization, we constructed strains lacking CDI systems (Δ*cdiBAI1*, Δ*cdiBAI2*), colicins (Δ*colBI*, Δ*colMI*, Δ*colYI*), all colicins (Δcolicins), or all competition systems (Δcomplete). These strains were competed against *E. coli* R12 or MG1655 targets lacking competition systems and immunity genes on solid media.

Wild-type R12 (WT) outcompeted Δ*cdiBAI1* and Δ*cdiBAI2* mutants by ∼10-fold (Fig. 2A), while controls (target cells provided with plasmid encoded CdiI immunity (p*cdiI*) or when the inhibitors lacked CDI systems (CDI^-^) showed no change.

**Figure 2.**
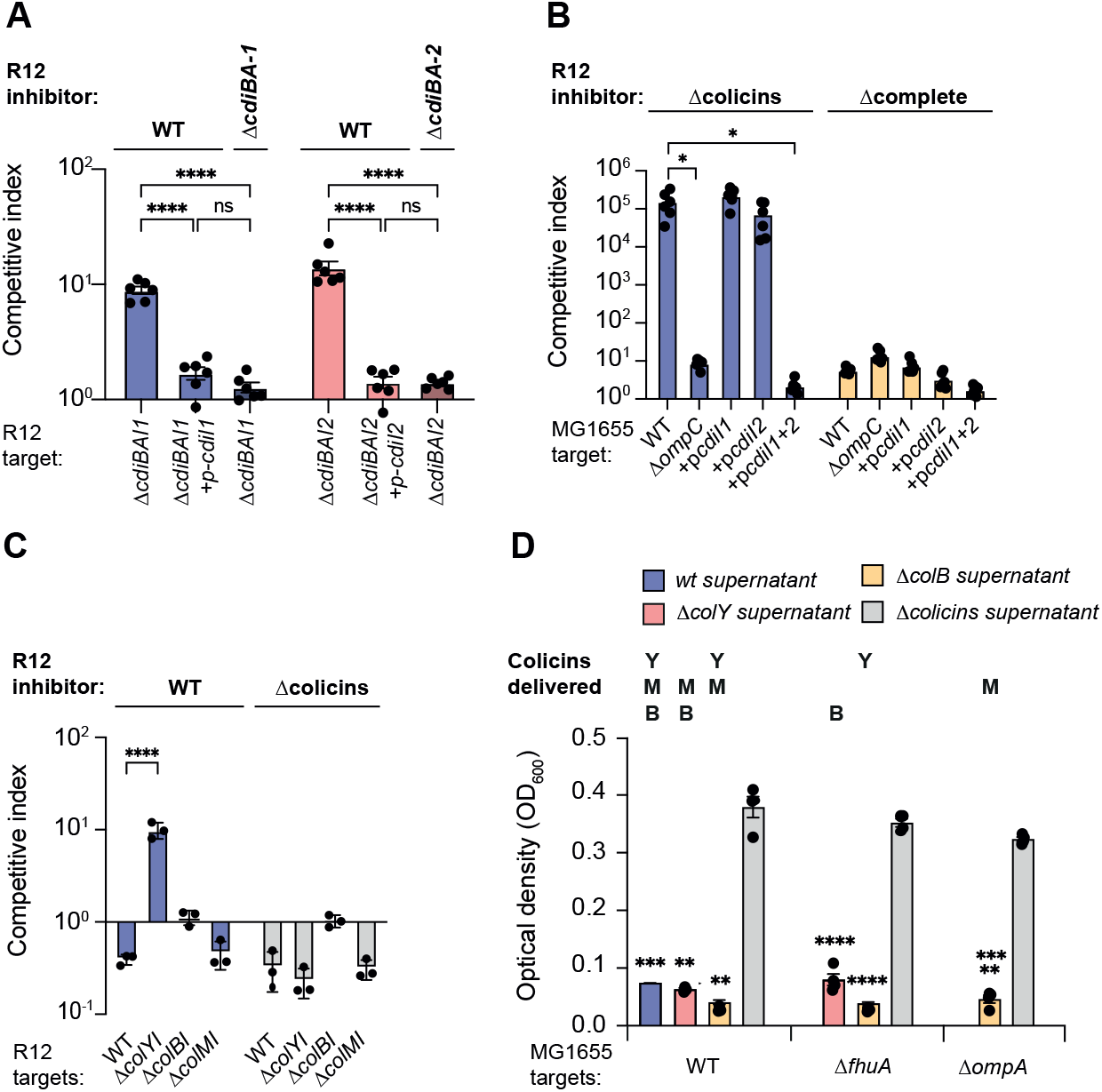
Colicin and CDI activity in *E. coli* R12. (A, B) To check for CDI activity, wild-type *E. coli* R12 and deletion mutants were co-cultured for 24 h on M9Gly+casAA solid media with (A) R12 lacking *cdiBAI1* or *cdiBAI2* or (B) MG1655 lacking *ompC* or complemented with CDI immunities. (C) Indicated R12 inhibitors were cocultured with R12 targets lacking *colYI, colBI* or *colMI* at a 10:1 ratio for 24 h on solid M9Gly+CAA minimal media to check for colicin activity. (D) MG1655 lacking the indicated colicin receptor were incubated with culture supernatant from R12 wt, Δ*colB*, Δ*colY* or Δcolicins for 4 h before measuring optical density. Data are the averages ± SEM of at least three independent experiments. For statistical analyses, two-way ANOVA with Tukey’s correction for multiple comparison (A, C), unpaired t-test (B, E) In D, significance was determined towards the Δcolicin supernatant for each target. ns, not significant, *, p ≤ 0.05, **, p ≤ 0.01, ***, p ≤ 0.001, **** ≤ 0.0001, ***** ≤ 0.00001.

R12 Δcolicins outcompeted MG1655 lacking single or double CDI immunity (p*cdiI1* or p*cdiI2* alone) by ∼4-logs (Fig. 2B). This advantage decreased to ∼1-log against Δ*ompC* targets (lacking CDI receptor) or strains expressing both *cdiIs* (p*cdiI1+ 2*)(Fig. 2B). The small competitive disadvantage of the Δ*ompC* strain is likely caused by a known fitness cost associated with deletion of *ompC* ^27^, as Δcomplete showed similar competition against MG1655 Δ*ompC*. Thus, both CDI systems are active and use OmpC as receptor.

To test colicin activity, R12 was competed against targets lacking individual immunity genes. Only Δ*colYI* targets were outcompeted ∼10-fold by WT R12, but not by R12Δcolicins; Δ*colBI* and Δ*colMI* targets were unaffected (Fig. 2C). These results suggest that colicin Y, but not colicins B or M, are active under these conditions.

As colicins are secreted to the extra-cellular milieu, we also assessed colicin activity in supernatants. MG1655 or immunity-deficient targets were exposed to overnight R12 culture supernatants. No inhibition occurred against R12 targets (Fig. S4A), but WT supernatant fully inhibited MG1655 (Fig. 2E). Using receptor knockouts (Δ*fhuA* colicin M, Δ*ompA* -colicin Y) and strains producing subsets of colicins, we assigned activity to each toxin. The colicin(s) produced and able to enter target is shown at the top of Fig. 2D. These combinations show that all three colicins are produced and active in supernatant, collectively inhibiting MG1655 growth.

There is a discrepancy in colicin-mediated growth inhibition between competition and supernatant assays. One possible explanation is the presence of full-length LPS on *E. coli* R12, which can block bacteriocins and antimicrobial peptides from accessing the cell surface via steric hindrance or charge-dependent sequestration ^47^. We find that MG1655 with full-length LPS ^48^ is fully resistant to R12 supernatant (Fig. S4A), as observed for R12. During competition, R12 outcompeted MG1655 with full-length LPS 10-fold (Fig. S4B), similar to the effect observed against R12 targets lacking colicin Y immunity (Fig. 2C). However, full-length LPS does not block CDI, as CDI-expressing R12 outcompetes MG1655 with full-length LPS, whereas colicin-expressing R12 does not (Fig. S4C). Taken together, colicins contribute to fitness, but target LPS structure modulates their effectiveness.

### Toxin delivery systems contribute to successful host colonization

To determine whether colicin and CDI toxin delivery systems contribute to colonization in a intact microbiota, we infected untreated mice with R12 lacking all five competition systems (Δcomplete) or WT R12 (Fig. 3A). Both strains showed similar colonization until day 6 p.i. but days 7-9 WT increased 10-to 100-fold, while Δcomplete did not (Fig. 3A). Thus, toxin delivery systems in R12 are critical for intestinal colonization.

**Figure 3.**
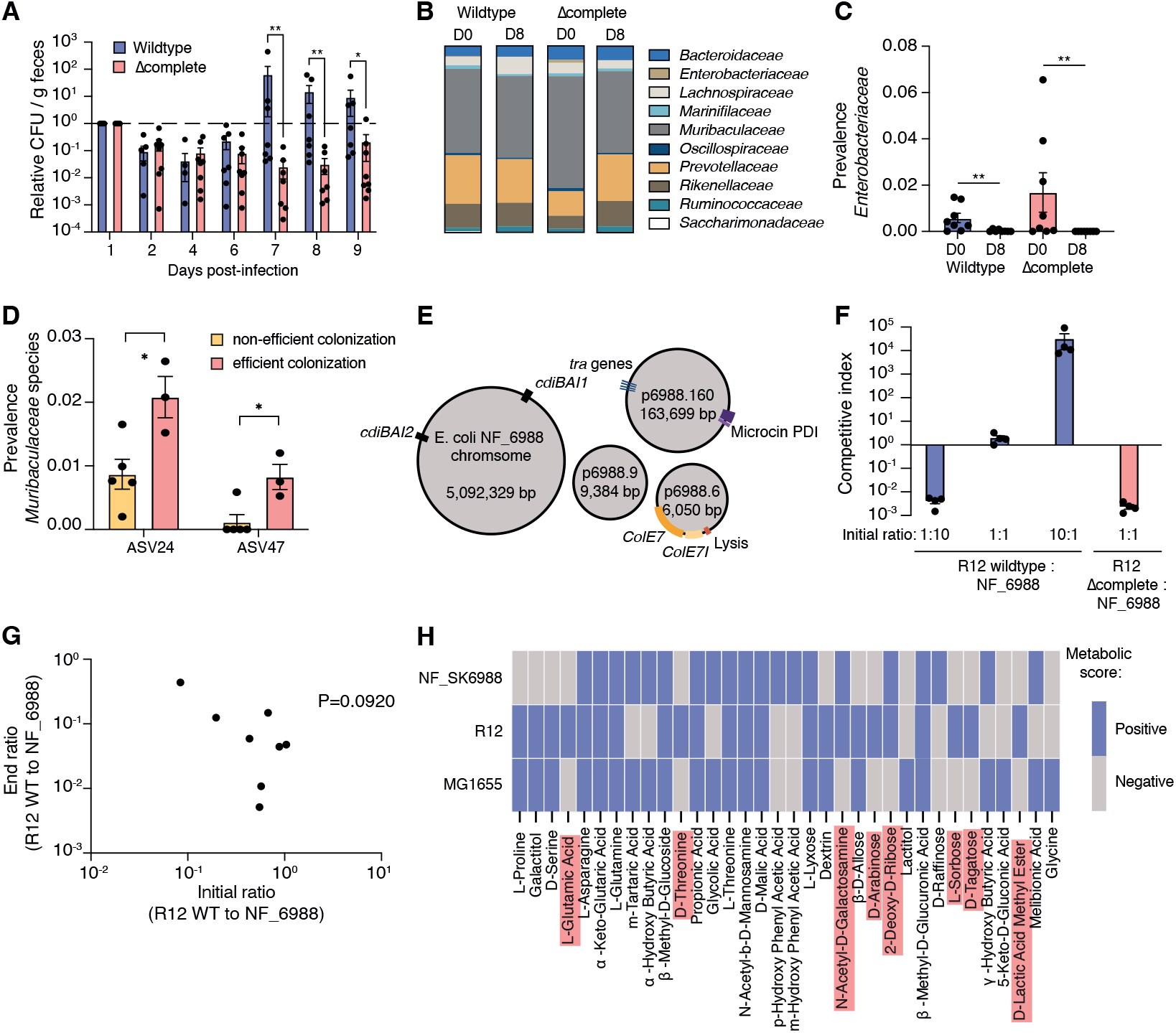
Colonization of untreated mice is improved by toxin delivery systems. (A) Untreated BALB/c mice were inoculated orally with 10^7^ wildtype or mutant R12 lacking all five toxin delivery systems (Δcomplete). The CFU levels were determined by plating fecal material on MacConkey agar plates with or without Rifampicin. The data is plotted as CFU/g feces relative to day 1. Each dot represents one mouse (N=7 wt and n=8 Δcomplete). P values were obtained using Mann-Whitney U test comparing the mutant strain to WT. *, p ≤ 0.05, ** p ≤ 0.01. (B) 16S sequencing results. (C) The prevalence of *Enterobacteriaceae* at D0 and D8 was compared for wildtype and Δcomplete using Mann-Whitney U test. ** p ≤ 0.01. (D) Statistically significant differences in *Muribaculaceae* species annotated as ASV24 and ASV47 in table S2 between mice colonized with R12 WT. Efficient colonization (n=3) is shown as yellow bars and non-efficient colonization (n=5) as pink bars.. Error-bars are SEM. Statistical significance was determined using unpaired t-test. * = P<0.05. (E) Overview of the *E. coli* NF_6988 genome. Genes encoding CDI systems, colicin E7, microcin PDI, and transfer proteins (*tra*) are indicated. (F) Coculture experiments of *E. coli* R12 wildtype and Δcomplete, and NF_6988 mixed at 10:1, 1:1 or 1:10 starting ratio on solid M9Gly+CAA minimal media for 24 h. (G) Correlation of the initial ratio and end ratio of R12 to resident *E. coli* microbiota was analyzed using Spearmans’s correlation. (H) The x-axis lists the 34 carbon sources in the Biolog PM1 and PM2 MicroPlates found to be different between MG1655, R12 and NF_6988. Carbon sources shaded in blue could be used by a strain as defined by a threshold of background-subtracted A_590nm_. Carbon sources utilized by R12 but not by NF_6988 are highlighted in bold and carbon sources utilized by R12 but not MG1655 are highlighted in red.

Microbiota composition changed during infection. *Enterobacteriaceae* decreased by day 8 in mice infected with both WT and Δcomplete, while the rest of the microbiota remained relatively stable (Figs. 3B–C, S6, Table S2). In Δcomplete-infected mice, *Muribaculaceae* increased, whereas *Prevotellaceae, Rikenellaceae*, and *Ruminococcaceae* decreased over eight days (Fig. S5). Some differences likely reflect initial microbiota variation, as Δcomplete mice had less *Muribaculaceae* and more *Prevotellaceae* at day 0 despite co-housing prior to the infection.

R12 colonization varied widely between mice for both WT and Δcomplete (Fig. 3A). Mice with efficient colonization had higher *Enterobacteriaceae* levels at the end of the experiment (Fig. S6).16S rDNA sequencing showed microbiota differences between mice, but no taxa differed significantly between efficient and inefficient colonization (Table S2). However, two *Muribaculaceae* species (ASV24 and ASV47) differed between these groups for R12 WT (Fig. 3D, Table S2). This analysis was not possible for Δcomplete, as efficient colonization occurred in only one mouse. Thus, R12 does not alter overall microbiota composition, but the initial microbiota outside *Enterobacteriaceae* influences *E. coli* colonization.

The toxin delivery systems in R12 are only known to target other *Enterobacteriaceae* ^20,27,49^. Consistent with this, *Enterobacteriaceae*, all of which were *E. coli*, decreased during the infection (Fig. 3C, Table S2). To further characterize this population, we analyzed OmpC variable loop sequences from 40 isolates collected before R12 gavage. A single OmpC-type was found in all mice, termed *E. coli* NF_6988 (NF – normal flora). Genome sequencing identified several competition systems; colicin E7, microcin PDI, and two *cdiBAI* loci in NF_6988 (Fig. 3E) (PRJNA1145732 / SAMN43077747). We next tested competitions between *E. coli* NF_6988 and R12 (WT or Δcomplete) at different starting ratios. At a 10:1 ratio, R12 outcompeted NF_6988 by about 5-logs (Fig. 3F). In comparison, at 1:10 ratio, NF_6988 outcompeted R12 by 2-3 logs. At a 1:1 ratio NF_6988 outcompeted R12 Δcomplete by ∼3-logs (Fig. 3F, pink bar). These results indicate that both initial ratio and toxin repertoire determine competitive outcome.

We next examined R12:NF_6988 ratios on day 1 of the mice infection. Ratios >10 occurred once for WT and twice for Δcomplete. This led to efficient colonization for WT, but not Δcomplete (Fig. S7). Most mice had ratios between 1:1 and 1:10, with little colonization (except one Δcomplete case). In two cases, ratios <1:10 still led to efficient colonization. However, no significant Spearman correlation was found between initial ratio and colonization success (Fig. 3G). Thus, while high initial ratios (>10:1) favor R12 colonization, other factors influence outcomes at lower ratios.

Other studies indicate that carbon nutrition influences bacterial growth in the mouse intestine ^7,50-52^. To compare carbon-utilization by R12 and the microbiota isolate NF_6988, we used Biolog PM1-2 MicroPlates, profiling 190 carbon sources. *E. coli* MG1655 was included as a reference. We identified 17 carbon metabolized only by one strain (Fig. 3G). Out of these, R12 utilized 11 not used by NF_6988. Unlike NF_6988, R12 can use galactitol as sole carbon source, a component of mouse chow shown to influence the co-existence of *E. coli* and *Salmonella* ^51,53^. It is possible that R12’s ability to use galactitol, or other unique carbon sources, may enable it to dominate the *E. coli* niche persist in the gut even in the absence of competitive systems.

### Colicins and CdiA toxins both contribute to colonization ability

The natural microbiota of mice retrieved from Charles River laboratories contained a competive *E. coli* isolate, requiring a high R12:NF ratio for R12 to dominate. To investigate how toxin delivery system affect colonization without resident competition, we used mice with a controlled microbiota (Fig. 4A). Given that *E. coli* R12 primarily impacts the *Enterobacteriaceae* niche, we pre-treated the mice with streptomycin to deplete *Enterobacteriaceae* before colonizing with *E. coli* MG1655 microbiota lacking known competition systems. Mice were then infected with wildtype or mutant R12 strains lacking CDI and/or colicins. By day 2, R12 WT comprised 90–100% of the *E. coli* population, and by day 6 it had eliminated MG1655 in all mice (Fig. 4B). In comparison, MG1655 could not displace an established MG1655 population (Fig. S8). Mutants lacking CDI (ΔCDI) showed bimodal colonization where R12 displaced MG1655 in some mice but remained low in others (Fig. 4B). Similarly, Δcolicins mutants showed impaired, bimodal colonization. The Δcomplete strain showed reduced early colonization but later reached WT levels (Fig. 4B). These results suggest that competition systems are important early in colonization, while other factors, such as nutrient use, support persistence. Using Biolog data (Fig. 3H), R12 was found to grow on seven carbon sources not used by MG1655, indicating that metabolic advantages may drive outgrowth independently of toxin delivery.

**Figure 4.**
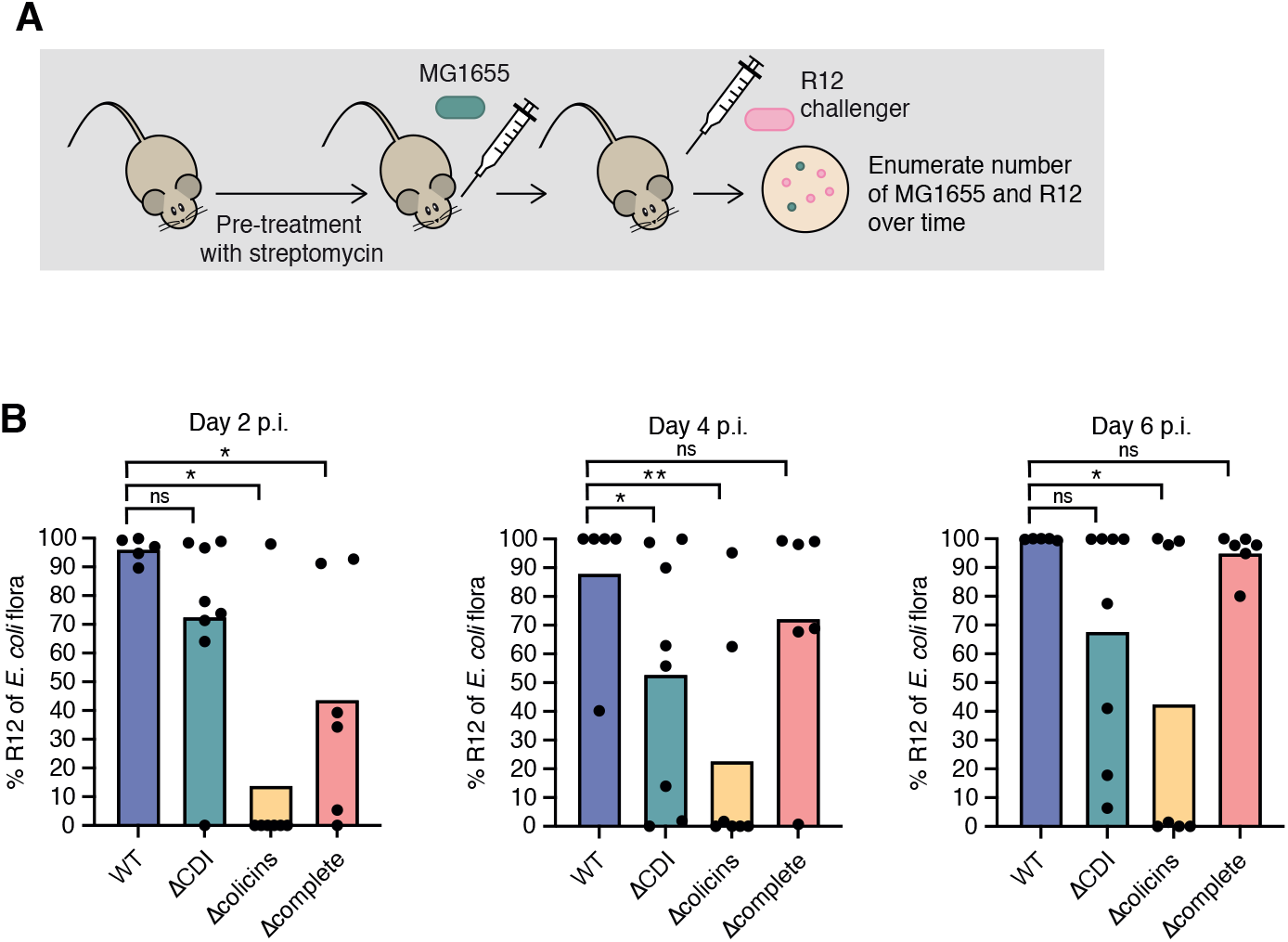
Colonization is dependent on CDI and colicins. (A) Schematic overview of mouse experiment. Antibiotic-treated BALB/c mice were orally inoculated with 10^7^ *E. coli* K12 MG1655 and subsequently challenged with 10^7^ of either wildtype or mutant *E. coli* R12. Each day fecal samples were collected and CFU counts were determined. (B) Mice pre-colonized with MG1655 were challenged with wildtype, or mutants lacking both CDI systems (ΔCDI), all three colicins (Δcolicins) or CDI systems and colicins (Δcomplete), and CFU counts of MG1655 and R12 were determined from fecal samples. Each dot represents one mouse. P values were obtained using Mann-Whitney U test comparing the mutant strains to WT. *, p ≤ 0.05.

We then tested whether bimodal colonization reflected differences in initial invader:resident ratios. A significant Spearman correlation between day 0 and day 6 ratios was observed for WT (P = 0.002232) and Δcolicins (P = 0.01667) (Fig. S9AB), but not for ΔCDI (P = 0.09618) or Δcomplete (P = 0.171) (Fig. S9CD). This supports that initial ratios influence colonization when competition systems are present. Overall, competition systems, nutrient utilization, and inoculum size shape colonization success, with the initial invader:resident ratio playing a key role in determining outcome.

## Discussion

The microbiota composition is important for our health, but detailed understanding of how bacteria invade occupied niches is lacking. Here we use *E. coli* as a proxy, to show that narrow-range bacterial weapons such as CDI systems and colicins are important in bacterial colonization of an intact niche, but that their efficacy largely depends on the toxin arsenal and metabolic abilities of the resident microbiota. In addition, we find that competitive *E.coli* can invade a niche occupied by resident bacteria with weapons of their own, but that this requires high invader:resident ratio at the initial colonization event. At lower ratios, or when the invader lacks toxin delivery systems, colonization efficacy becomes more erratic and susceptible to other factors.

Previous work indicates that metabolic capabilities influence gastrointestinal colonization success ^7,50,51^. We find that R12 can utilize several rare carbon sources, some important in the gut and others not metabolized by competing *Enterobacteriaceae*. Variation in colonization efficiency may therefore reflect differences in non-*Enterobacteriaceae* microbiota, affecting the availability of these metabolites in the gut. Notably, two *Muribaculaceae* species (ASV24 and ASV47) differ between mice with efficient versus poor R12 WT colonization. *Muribaculaceae* metabolize polysaccharides including plant and host glycans, producing monosaccharides that potentially fuel other resident microbiota ^54^. However, metabolic capacity varies across species within the family ^55^. For example, *Muribaculaceae gordoncarteri* show high a-arabinase and a-fucosidase activity ^56,57^, whereas *Heminiphilus faecis* ferments mannan and L-arabinose ^58^. Thus, it is possible that these particular species of *Muribaculaceae* produce certain metabolites important for R12 colonization. Further work is needed to determine how strain-specific metabolism shapes competition and colonization.

On the other hand, *Muribaculaceae* also produce short-chain fatty acids and regulate the intestinal barrier function and the immune response ^59^. The CdiA protein contains multiple filamentous haemagglutinin (FHA) repeats that both induce production of, and are recognized by, secretory IgA (sIgA). ^60-62^. sIgA recognition of bacterial surface molecules results in clumping of the cells, prevention of motility and clearance from the gut via intestinal fluid flow ^63,64^. The resident *E.coli* NF_6988 encoded similar CdiA proteins as R12. Thus, it is possible that in mice with high levels of these particular *Muribaculaceae* have a different immune response (e.g. lower levels of CdiA responsive sIgA), resulting in improved colonization efficacy of R12. This is further supported by our finding that R12 Δcomplete colonizes mice with MG1655 microbiota more effectively than R12 Δcolicins. If CDI-expressing cells (R12 Δcolicins) are more susceptible to sIgA-mediated clumping and clearance than ΔCDI strains, which lack FHA domains recognized by sIgA, this could explain their reduced colonization efficiency relative to Δcomplete. As the prevalence of specific *E. coli* strains in the microbiota varies between individuals and over time ^65^, our susceptibility to specific *E.coli* strains might also vary.

Strain-level variation in *Enterobacteriaceae* matters because different *E. coli* strains cause diseases ranging from diarrhea and urinary tract infections to cancer and inflammatory bowel disease ^3,66,67^. Although little is known about such variation in healthy gut microbiomes, travel studies show frequent colonization by antibiotic-resistant strains in high-AMR regions ^68^. Most travelers lose these strains after returning, but some retain them for years ^69^. Our data suggest colonization success depends on growth competition and nutrient use of invading and resident strains. Narrow-range antagonistic systems in *E. coli* R12 target *Enterobacteriaceae*, offering potential for targeted microbiota modulation, such as removing pathogenic or resistant strains. Overall, this work highlights molecular bacterial interactions in shaping strain-level variation and guiding microbiota interventions.

## Material and methods

### Strains and growth conditions

The bacterial strains used in this study are listed in table S3. *E.coli* R12 was isolated from the feces of rats acquired from Charles River laboratories. All strains were grown at 37°C and with shaking at 200 rpm. in M9Gly+CAA minimal media: 1x M9 salts supplemented with 1% glycerol, 0.2% cas-amino acids, 2 mM MgSO_4_ and 0.1 mM CaCl_2_, or in Lysogeny broth (LB): 10 g l^-1^ tryptone, 5 g l^-1^ yeast extract and 10 g l^-1^ NaCl. Agar was added at 15 g l^-1^ for solid media. Media were supplemented with antibiotics when applicable as follows: ampicillin 100 mg l^-1^, chloramphenicol 12.5 mg l^-1^, kanamycin 50 mg l^-1^, streptomycin 100 mg l^-1^, rifampicin 100 mg l^-1^.

### Whole-genome sequencing

Chromosomal DNA from *E. coli* R12 and NF_6988 were isolated using the Qiagen Genomic-tip 100/G kit according to the manufacturer. PacBio libraries were produced using the SMRTbell™ Template Prep Kit 1.0 according to manufacturer’s instructions (Pacific Biosciences, CA, USA). Libraries were subjected to exo treatment and PB AMPure bead wash procedures for clean-up before size selection with the BluePippin system (Sage Sciences, MA, USA) with a cut-off value of 6500 bp. The libraries were sequenced on the PacBio RS II instrument using C4 chemistry, P6 polymerase and 240-minute movie time in one SMRTcell™. The Pacbio reads were assembled using HGAP3 from SMRTportal (PacBio, CA, USA) with default settings.

For NF_6988, barcoding and library preparation were performed using the Native Barcoding Kit 24 V14, followed by sequencing on an R10.4 flow cell in a MinION Mk1c device (Oxford Nanopore, Oxford, UK). Base calling and adapter trimming were performed with Guppy version 6.3.8+d9e0f64 (Oxford Nanopore) using the super-accuracy configuration (dna_r10.4.1_e8.2_260bps_sup). The resulting reads were then assembled using Flye v. 2.9.2-b1786 ^70^ and inspected using Bandage ^71^. Long-read polishing was performed using Medaka v. 1.0.3 (https://github.com/nanoporetech/medaka). Both genomes were then annotated with Prokka 1.14.6 ^72^ before being submitted as complete genomes to NCBI.

### Colicin production assay

Bacterial overnight cultures were centrifuged at 3,000xg for 5 minutes and the culture medium supernatant was passed through a 0.2 µm filter to make spent media. Overnight cultures of test strains were diluted 1:100 in M9 minimal medium, mixed with sterile-filtered spent medium at 1:25 ratio and incubated for 4 h at 37°C before measuring the optical density at 600 nm. Spent media were prepared from *E. coli* R12 wildtype, Δcolicin Y, and Δcolicins. Test strains were *E. coli* R12 mutants lacking the colicins and corresponding immunities, or *E. coli* K12 MG1655 lacking *fepA, fhuA, ompA*, alone or in combinations.

### Competition assay

Overnight cultures of inhibitor and target cells were mixed at a ratio of 10:1 and 20 µl were spotted onto solid M9Gly+CAA minimal media. The cells were co-cultured for 24 h at 37°C before suspended in 1xPBS and plated onto LB solid media containing appropriate antibiotics to enumerate the number of colony-forming units per milliliter of inhibitors and targets. Competitive indices were calculated as the ratio of inhibitors to target cells at the end of the co-culture divided by the ratio at the start of the co-culture.

### Animal experiments

Forty-eight 7-week old female Balb/c (Charles River laboratories distributed by Scanbur) were housed in individual ventilated cages (Tecniplast boxunsfue) at the Swedish National Veterinary Institute. The mice were housed 8 mice per cage in a controlled environment (23°C ± 2, 75 ACH and 12h light and dark cycle). The mice’s diet was given an autoclaved pellet feed with no less than 18% protein, 5% fat and 5% fibre, tap water and the cages was enriched with autoclaved hay, a house and a tunnel. All personal handling the mice were required to change clothes, shoes and wear overall, gloves and cap before entering the room, also gloves were changed between cages. All the mice were handled with non-aversive tunnel handling since it has been shown to reduce stress and anxiety in laboratory mice ^73^. After a seven-day acclamation period the mice were split into 4 mice per cage before the start of the experiment. For all inoculums, a single colony of each bacterial strain was grown in LB overnight. The gavage solution was prepared by diluting the overnight culture 1:10 in 1x Phosphate Buffered Saline (PBS) with bicarbonate solution.

For the experiment with undisturbed microbiota, mice were orally gavaged with 100 μl *E. coli* R12 wildtype or Δcomplete (10^7^ CFU/mouse). To ensure the bacteria did not affect the mice in any negative way the mice were closely watched after treatment and weighed every day to ensure no significant amount of weight were lost. Fecal samples were taken for CFU counts on days -5, -2, 0-9 as well as DNA extraction and metagenomic sequencing on days 0 and 8.

For the pre-colonization experiment, mice were given 5 g l^-1^ streptomycin in drinking water for 7 days. At day -2 p.i., mice were treated with 20 mg streptomycin per mouse by gavage. At day -1 p.i. streptomycin was removed from the water and mice were orally gavaged with *E. coli* K12 MG1655 (10^7^ colony-forming units (CFU)). The following day, challenger bacteria, *E. coli* R12 wildtype, ΔCDI, Δcolicins or Δcomplete were introduced by oral gavage (10^7^ CFU). Fecal samples were taken for CFU counts on days -5, -2, 0-9.

To determine the colony counts, each day fecal samples were obtained from each mouse. Fecal samples were weighed and resuspended in PBS using a plunger. The samples were serially diluted, plated on MacConkey agar supplemented with appropriate antibiotics, and CFU were determined by colony counting.

### Biolog PM metabolic phenotype analysis

The phenotypic analysis of *E. coli* MG1655, *E. coli* R12 and *E. coli* NF_6988 were conducted using Biolog Phenotype Microarray (PM) plates, PM1 and PM2, following the recommended protocol (Biolog Inc., Hayward, CA, USA). In summary, the bacteria were inoculated in LB liquid media and incubated for 16 h prior to diluting the sample 1:35 in IF-0 to 42%T (OD_600_

∼0.17) and subsequently diluted in IF-0+Dye to 85%T (OD_600_ ∼0.03). 100 µl per well were loaded on PM1 and P2 metabolic plates and the plates were incubated at 37°C, with orbital shaking, in an Infinite M200 microplate reader (Tecan). Absorbance at 590 nm was measured every 15 minutes for 48 h.

## Ethics

This study was conducted in accordance with Swedish animal welfare laws. All procedures were done according to LASAs good practice guidelines and handling was done with non-aversive tunnel handling to improve animal welfare ^73^. The Uppsala Animal Experiments Ethics review board in Uppsala, Sweden approved all mice protocols undertaken in this study under reference no. 5.8.18-5552/2019.

## Data Availability Statement

All raw data for the manuscript are freely available either as supplementary excel files (Tables S2 and S3) or at the NCBI data repository for genome sequences (accession numbers are found in the text).

## Author Contributions

P.M., D.L., and S.K. conceived the study. P.M., E.K., and S. K. performed experiments. P.M., J.K, E.K., and S.K analyzed data. P.M., D.L., and S.K. wrote the manuscript.

## Acknowledgements

This study was funded by grants from the Swedish research council, the foundation to prevent antibiotic resistance and Uppsala antibiotics center to S.K. We thank Mia Phillipson and Susan Schlegel for insightful input during the writing of this manuscript.

