## Supplementary material for "Roles of bacterial growth competition systems in colonization of the murine gut"

### Supplemental material for: Roles of bacterial growth competition systems in colonization of the murine gut.

**This file includes:**

**Supplementary methods**

**Supplementary tables 1-6**

**Supplementary figures 1-9**

**Supplementary references**

### Supplementary methods:

**Chromosomal and plasmid constructs**

Plasmids and oligonucleotides used in this study are listen in tables S5-S6.

The *cdiBA-1* and *cdiBA-2* were deleted using lambda red recombineering where the *cdiA* ORFs first was replaced by a chloramphenicol resistance maker amplified from pKD3 using oligos 1604+1605 and 1575+1576 for *cdiA1* and *cdiA2* respectively. The *cdiI1* and *cdiI2* immunity genes were subsequently removed in these strains using kanamycin-cassettes amplified from pKD4 with oligos 1604+1606 and 1575+1577. Verification was made using oligos 1607+1608 and 1607+1581 for CDI locus 1 and oligos 1578+1579 and 1582+1583 for locus 2. The resistance genes were removed using pcp20 mediated FLP-recombination as described previously [1].

To construct plasmids expressing constitutive immunity, *cdiI-2* was amplified from *E. coli* R12 genomic DNA using oligos 1856+1857 and cloned onto pTrc99aKX using KpnI and XhoI. Verification was made with oligos 2508+2509. This plasmid and pTrc99aKX::*cdiI-1* [2] were introduced into *E. coli* R12 *cdiBAI* KO strains.

*cdiI-2* was also amplified from E. coli R12 using oligos 2538+2771 and cloned into pSC101 under the control of pJ23101 promoter using BamHI and HindIII. Verification was made using oligos 488+483. Plasmids pTrc99aKX::*cdiI-1* [2] and pSC101-pJ23101::*cdiI-2* were introduced into *E. coli* K12 MG1655 *rpsL*(K42R).

Colicin B, colicin M and colicin Y and their immunities were deleted in R12 wildtype and Δ*cdiBAI-1* Δ*cdiBAI-2* strains using lambda red recombineering. KO cassettes were amplified from pKD3 using 1845+1846 (*colM*) and 1845+2128 (*colMI*), 1841+1842 (*colB*) and 1842+2127 (*colBI*), and 1613+1614 (*colY*) and 1613+2105 (*colYI*). Verification was made using primers 2129+1847 and 1920+1848 (*colMI*), 2130+1921 and 1843+1844 (*colBI*), and 1615+1616 and 2106+2107 (*colYI*).

Rifampicin resistance was introduced in R12 using Lambda red of *rpoB*(D516V), and verified by sequencing using primers 1879+1880.

Receptor deletion mutants were transferred from the Keio collection [3] of *E. coli* single-gene knockout strains using bacteriophage P1 mediated transduction. *ompC::kan* was introduced in *E. coli* K12 MG1655 *rpsL*(K42R) for CDI competitions. *fepA::kan*, *fhuA::kan* and *ompA::kan* were introduced into *E. coli* MG1655 *lacA-cat*. In order to make the double receptor deletion mutants, the kanamycin-cassette from the first deletion was removed using pCP20 encoded FLP recombinase before introducing the second deletion.

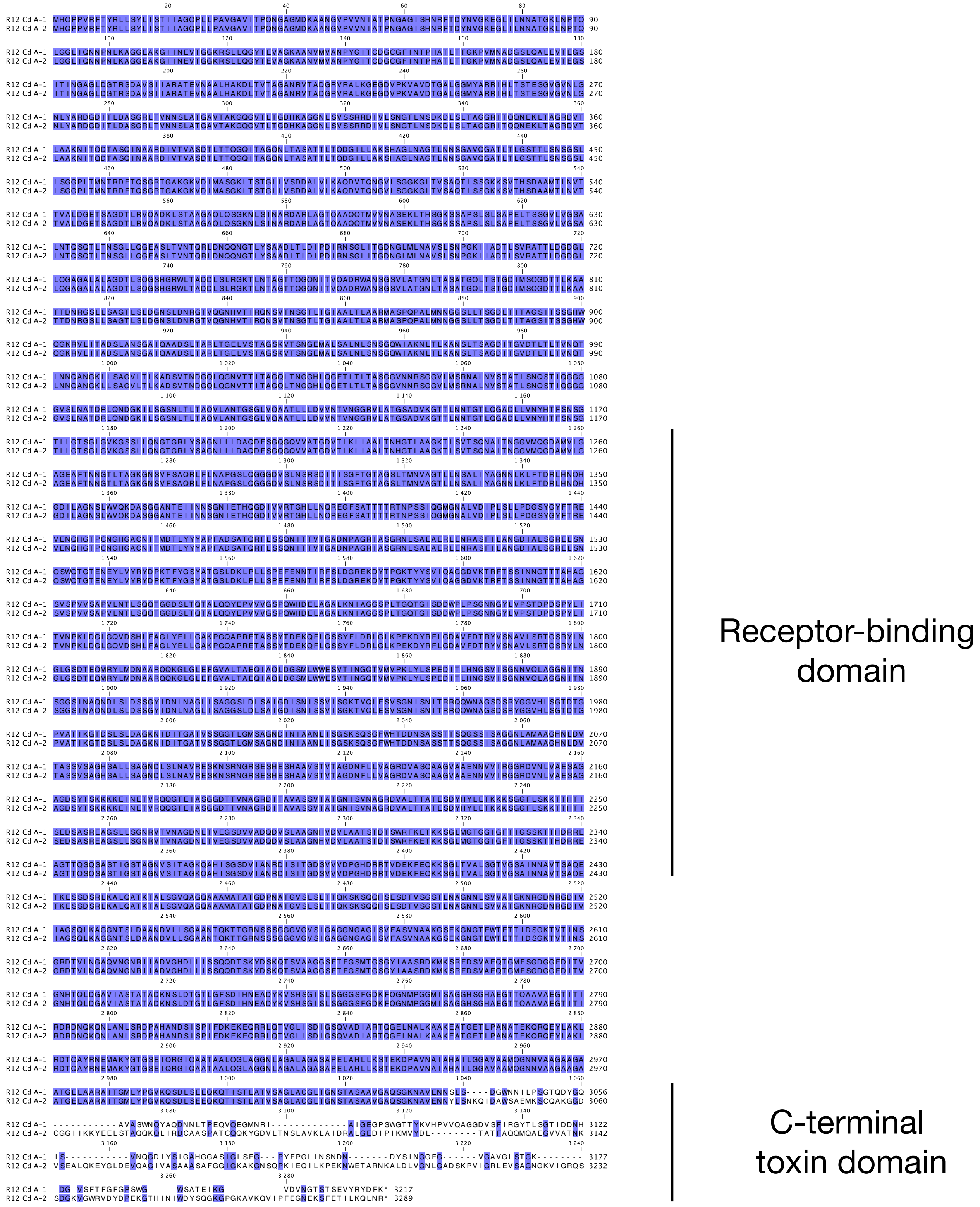


**Figure S1. Alignment of *E. coli* R12 CdiA-1 and CdiA-2.** The receptor-binding domain and C-terminal toxin domain are indicated. Identical residues are colored in blue, differences are indicated as lack of color.

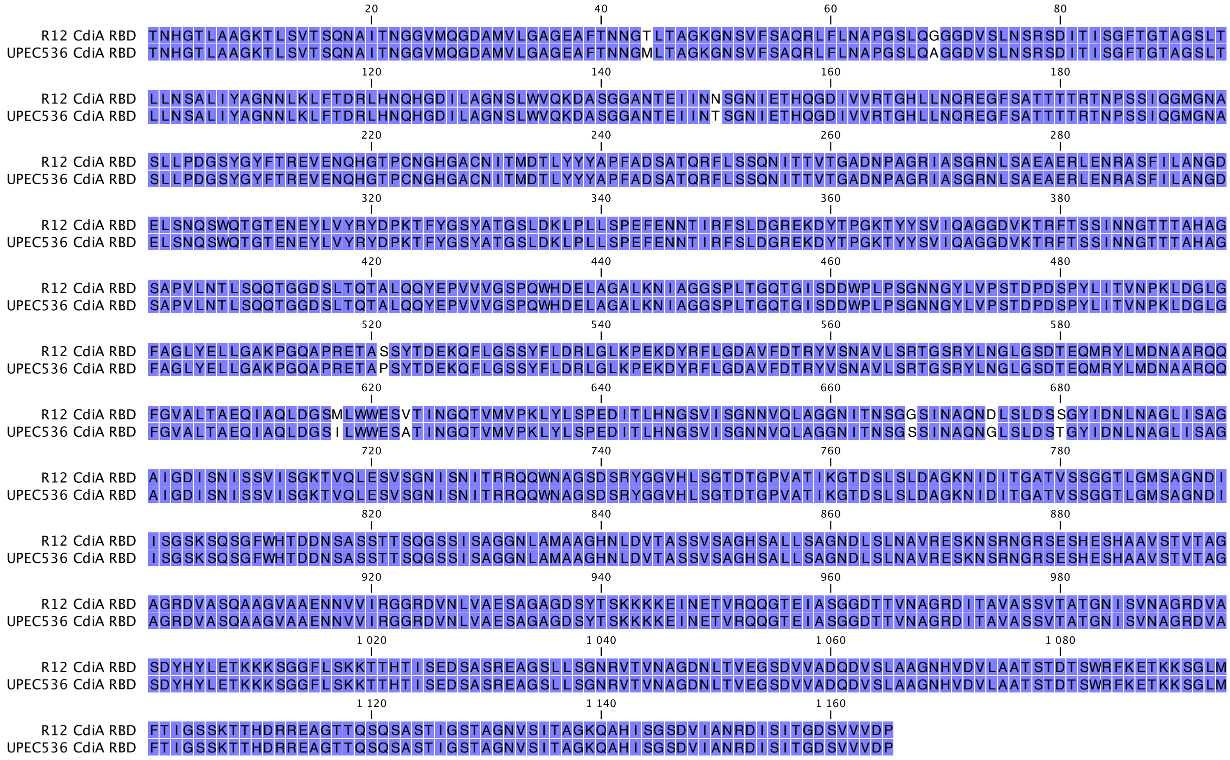


**Figure S2. Alignment of the CdiA receptor-binding domains from *E. coli* R12 and UPEC536.** Identical residues are colored in blue, differences are indicated as lack of color.

```
  R12 CdiA-CT1 VENNSLSDGWNNILPSGTQDYGQAVASWNQYAQDNNLTPEQVQEGMNRIAIGEGPSWGTTYKVHP 65
 EC93 CdiA-CT2 VENNSLSDGWNNILPSGTQDYGQAVASWNQYAQDNNLTPEQVQEGMNRIAIGEGPSWGTTYKVHP 65

  R12 CdiA-CT1 VVQAGGDVSFIRGYTLSGTIDDNHISVNQGDIYSIGAHGGASIGLSFGPYFPGLINSNDNDYSIN 130
 EC93 CdiA-CT2 VVQAGGDVSFIRGYTLSGTIDDNHISVNQGDIYSIGAHGGASIGLSFGPYFPGLINSNDNDYSIN 130

  R12 CdiA-CT1 GGFGVGAVGLSTGKDGVSFTFGFGPSWGWSATEIKGVDVNGTSTSEVYRYDFK 183
 EC93 CdiA-CT2 GGFGVGAVGLSTGKDGVSFTFGFGPSWGWSATEIKGVDVNGTSTSEVYRYDFK 183
```

**Figure S3. Alignment of C-terminal toxin domains from *E. coli* R12 CdiA-CT1 and *E. coli* EC93 CdiA-CT2.** Identical residues are colored in blue, differences are indicated as lack of color.

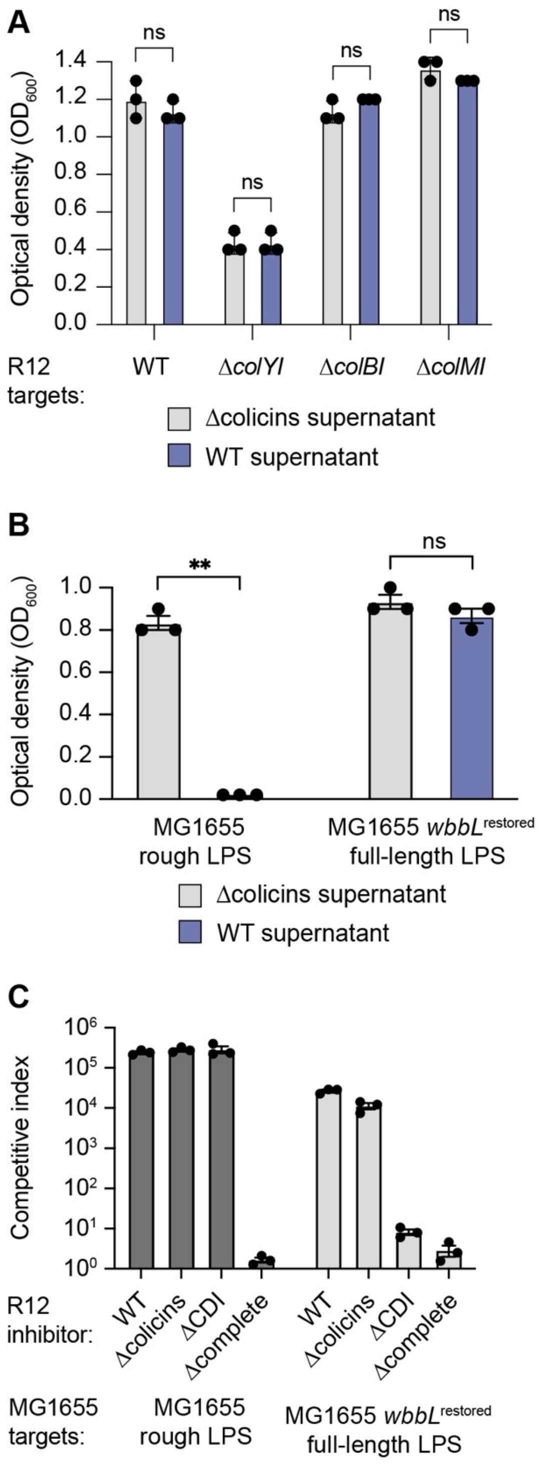


**Figure S4. R12 colicins are blocked by LPS.** Indicated R12 (A) or MG1655 (B) strains were exposed to spent media from R12 Δcolicins (grey) or WT (blue) and grown in M9Gly+CAA minimal media for 4 h before measuring optical density. (C) Indicated R12 inhibitors were cocultured with MG1655 wildtype (dark grey bars) or *wbbL*$^{restored}$ (light grey bars) at a 10:1 ratio for 24 h on solid M9Gly+CAA minimal media. Data are the averages ± SEM of three independent experiments. For statistical analyses, unpaired t-test were performed. ns, not significant, *, p ≤ 0.05, **, p ≤ 0.01.

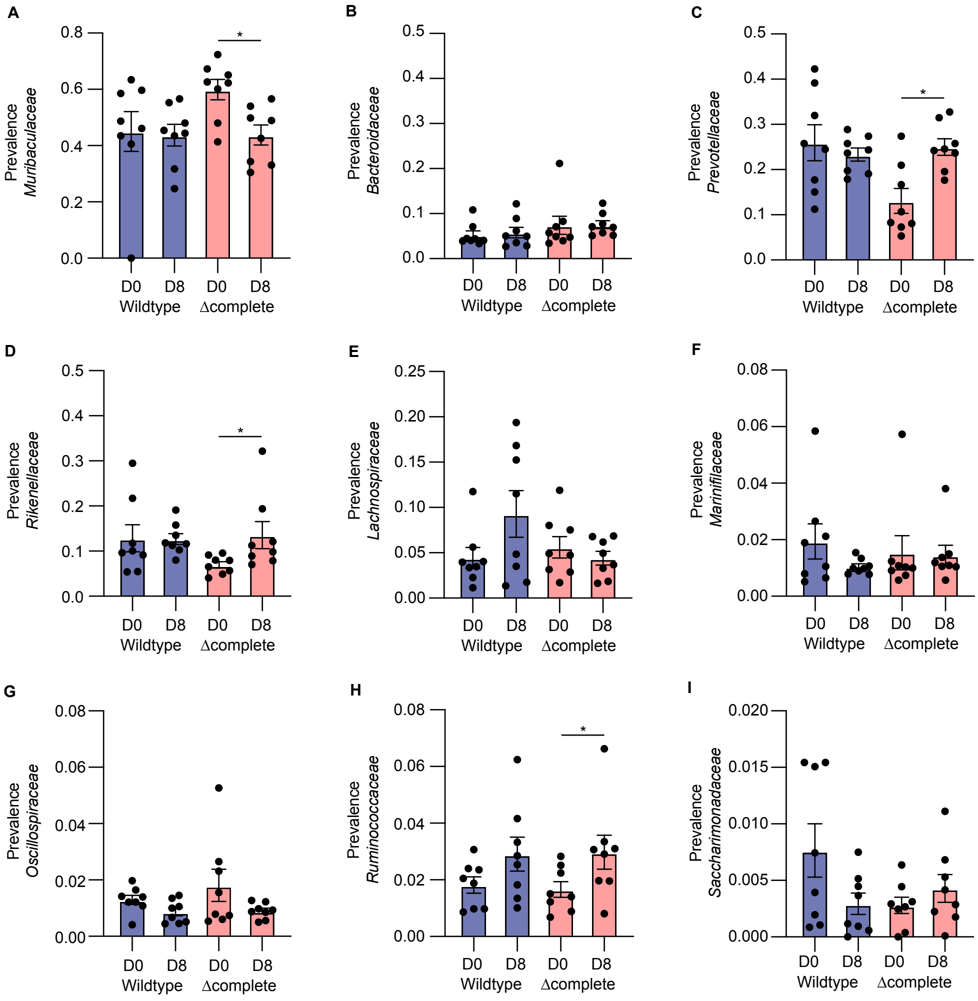


**Figure S5. Prevalence of different bacterial species in the mice microbiota**. The prevalence of (A) *Muribaculaceae*, (B) *Bacteroidaceae*, (C) *Prevotellaceae*, (D) *Rikenellaceae*, (E) *Lachnospiraceae*, (F) *Marinifilaceae,* (G) *Oscillospiracae*, (H) *Ruminococcaceae* and (I) *Saccharimonadaceae* at D0 and D8 was determined by 16S rDNA sequencing for wildtype and Δcomplete. Statistical analysis was done using Mann-Whitney U test. * $p \leq 0.05$.

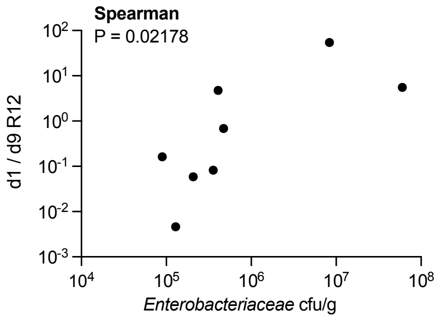


**Figure S6. Correlation between ability to cause efficient colonization and final enterobacterial load.** Spearman correlation of ability to cause efficient colonization for R12 wt and the final enterobacterial load in cfu/g.

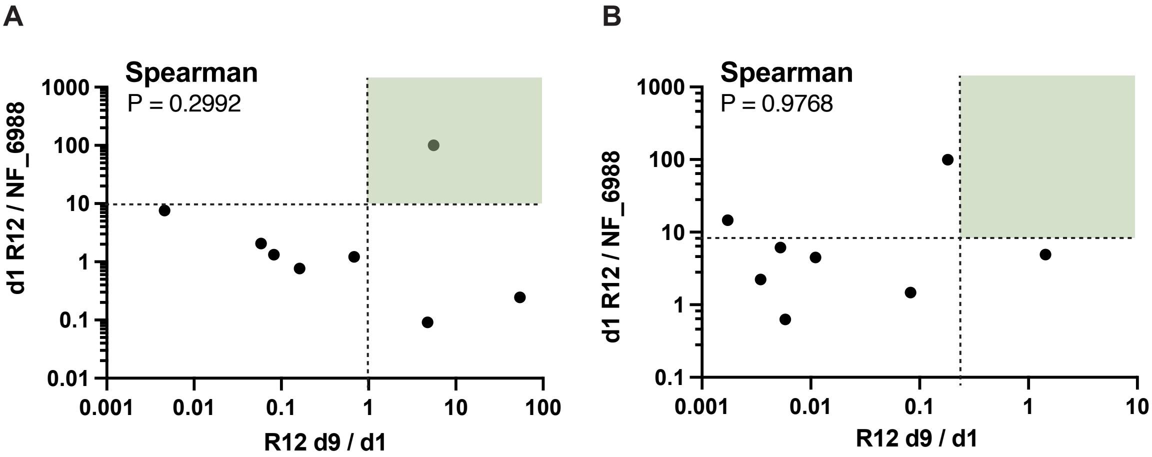


**Figure S7. Correlation between initial ratio of R12 and NF_6988 and the ability to cause efficient colonization.** Spearman correlation of initial (d1 y-axis) ratio of R12 to NF_6988 and ability to cause efficient colonization for wt (A) and Δcomplete (B). Efficient colonization was determined as an increase in prevalence (d9/d1 ratio > 1) indicated by a vertical hatched line. Initial ratio of R12:NF6988 at 10:1is demarcated by a horizontal hatched line. The zone with high colonization efficiency and starting ratio > 10:1 is indicated in green.

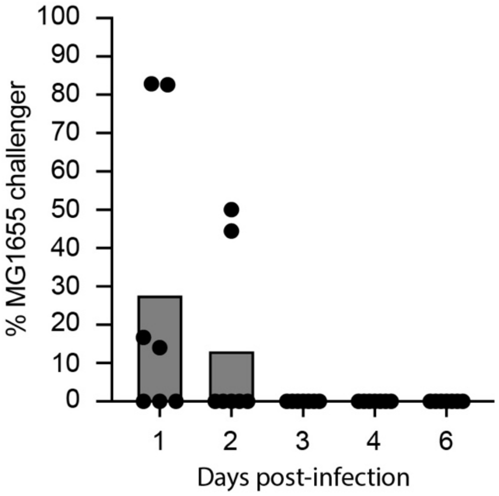


**Figure S8**. **MG1655 colonization.** Mice pre-colonized with MG1655 were challenged with MG1655 and CFU counts of both strains were determined from fecal samples at indicated days. Each dot represents one mouse.

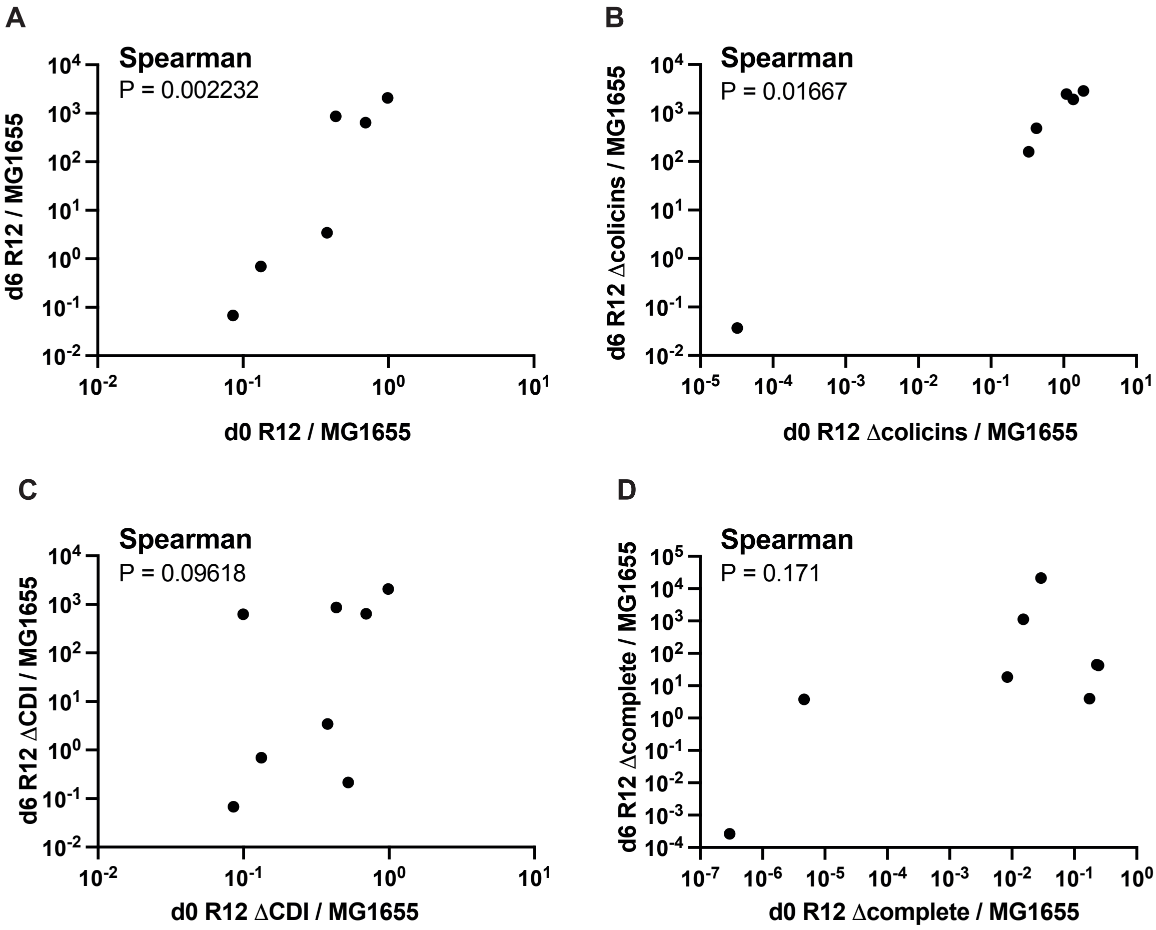


**Figure S9. Correlation between initial and end ratio of R12 to MG1655.** Spearman correlation of initial (d0 x-axis) and end (d6 y-axis) ratio of *E.coli* R12 vs. *E.coli* MG1655. (A) R12 wt, (B) Δcolicins (C) ΔCDI and (D) Δcomplete.

**Table S1. Genomic comparison of *E. coli* R12 and E. coli MP1.** Sequence comparison between chromosome and different plasmids found in *E.coli* R12 and MP1 performed with pyani v. 0.2.12. Similarity between two sequences is calculated as the proportion of nucleotide identity multiplied with the proportion of alignment coverage. Thus, the maximum similarity of 1.0 represents plasmid sequences that are 100% identical and completely overlap with each other. The comparison of R12 and MP1 genomes shows that they are not the same strain, even though they share 98% similarity on the chromosome and 87% on the pColY plasmid, R12 also contains the *colBM* plasmid, not present in MP1.

| | **R12 pColY_like plasmid** | **MP1 chromosome** | **MP1 plasmid** | **R12 chromosome** | **R12 81kb plasmid** |
|---|---|---|---|---|---|
| **R12 pColY_like_plasmid** | 1.0 | 0.0 | 0.87 | 0.0 | 0.0 |
| **MP1 chromosome** | 0.0 | 1.0 | 0.0 | 0.99 | 0.00018 |
| **MP1 plasmid** | 0.88 | 0.0 | 1.0 | 0.0 | 0.0 |
| **R12 chromosome** | 0.0 | 0.98 | 0.0 | 1.0 | 0.00037 |
| **R12 81kb plasmid** | 0.0 | 0.0099 | 0.0 | 0.022 | 1.0 |

**Table S2. Data from metagenomics experiment.**

Table S2 is found as a separate file.

**Table S3. Raw data for experiments included in the paper.**

Table S3 is found as a separate excel file and contains colony counts and processing of all competition and colonization experiments.

**Table S4. Bacterial strains used in this study**

| Strain number | Genotype | Origin |
|---|---|---|
| SK620 | *E. coli* K12 MG1655 *lacA-cat* | [4] |
| SK2743 | *E. coli* K12 MG1655 *araB::FRT-kanR-FRT rpsL(K42R)* | This study |
| SK3573 | *E. coli* R12 | D. Low |
| SK3935 | *E. coli* K12 MG1655 *rpsL(K42R)* | This study |
| SK4352 | *E. coli* R12 *cdiBA-2::cat* | This study |
| SK4395 | *E. coli* R12 *cdiBAI-2::kan* | This study |
| SK4524 | E. coli R12 *cdiBA-1::cat* | This study |
| SK4576 | *E. coli* R12 *cdiBAI-1::kan* | This study |
| SK5233 | *E. coli* R12 *rpoB(D516V)* | This study |
| SK5594 | *E. coli* R12 Δ*colY* | This study |
| SK5596 | *E. coli* R12 Δ*colY*, *cma::kan*, *cba::cat* | This study |
| SK5654 | *E. coli* R12 *colYI::cat rpoB(D516V)* | This study |
| SK5664 | *E. coli* R12 Δ*colY*, *cba::cat*, *cma::kan*, *rpoB(D516V)* | This study |
| SK5699 | *E. coli* R12 Δ*cma*, *cmi::cat*, *rpoB(D516V)* | This study |
| SK5700 | *E. coli* R12 Δ*cba*, *cbi::kan*, *rpoB(D516V)* | This study |
| SK5920 | *E. coli* R12 Δ*cdiBA-2* Δ*cdiBA-1* Δ*colY cbi-cmi::cat*, *rpoB(D516V)* | This study |
| SK6555 | *E. coli* R12 *rpoB(D516V)* | This study |
| SK6557 | *E. coli* R12 Δ*cdiBA-2* Δ*cdiBA-1 rpoB(D516V)* | This study |
| SK6620 | *E. coli* K12 MG1655 *lacA-cat, fepA::kan* | This study |
| SK6621 | *E. coli* K12 MG1655 *lacA-cat fhuA::kan* | This study |
| SK6622 | *E. coli* K12 MG1655 *lacA-cat ompA::kan* | This study |
| SK6623 | *E. coli* K12 MG1655 *rpsL(K42R) ompC::kan* | This study |
| SK6624 | *E. coli* K12 MG1655 *rpsL(K42R)* /pTrc99aKX*::cdiI-2*$^{EC93}$ | This study |
| SK6646 | *E. coli* R12 *rstA(Thr79) rpoB(D516V)* | This study |
| SK6791 | *E. coli* R12 Δ*cdiBA-2* Δ*cdiBA-1* Δ*colicinY colicinBI+MI::cat*, *rpoB(D516V)*, *rstA*(SK3574) | This study |
| SK6864 | *E. coli* K12 MG1655 *lacA-cat* Δ*ompA fepA::kan* | This study |
| SK6865 | *E. coli* K12 MG1655 *lacA-cat* Δ*ompA fhuA::kan* | This study |

| | | |
|---|---|---|
| SK6993 | *E. coli* K12 MG1655 *rpsL(K42R)* /pSC101-pJ23101::*cdiI-2* | This study |
| SK6995 | *E. coli* K12 MG1655 *rpsL(K42R)* pTrc99aKX:: *cdi-2*$^{EC93}$ /pSC101-pJ23101::*cdiI-2*$^{R12}$ | This study |

**Table S5. Plasmids used in this study.**

| **Plasmid** | **Genotype** | **Origin** |
|---|---|---|
| pSK2964 | pTrc99aKX:: *cdiI-2*$^{EC93}$ | [2] |
| pSK6553 | pTrc99aKX:: *cdiI-2*$^{R12}$ | This study |
| pSK6991 | pSC101-pJ23101::*cdiI-2*$^{R12}$ | This study |

**Table S6. Oligos used in this study.**

| Oligo | Sequence (5' to 3') |
|---|---|
| 483 | CCAGAACAGCCCGTTTGC |
| 488 | TTCCCGTTTCAGTGACAACG |
| 1575 | ATTTCAGCACTGACATTCCGCTTACGTTAATTTACACTGAtgtaggctggagctgcttc |
| 1576 | TCAAAAGACTTTTCATTGCCTTCAAAAGGTATGACTTGCTccatatgaatatcctccttagttcc |
| 1577 | CTATTTCTTGATTCCTAAACGGAACATATCAGTTGGGAATccatatgaatatcctccttagttcc |
| 1578 | AGTGGATTATGCTGCTATAGC |
| 1579 | TTGCGTTGTTACCCTGTC |
| 1581 | CATTGAGTCCCGCATGACTT |
| 1582 | GCTAAACAGGTAGAGCAA |
| 1583 | AGCTGTTACGTCATAGATATT |
| 1604 | TTCTGGTTGTTTCATCCCCGCATTTTCGTCTCCGGCAGCCtgtaggctggagctgcttc |
| 1605 | ATCGATAAACCTCACTTGTTGAAGTGCCATTAACGTCAACccatatgaatatcctccttagttcc |
| 1606 | CATATTCACTACCGCCAAGTAAATTATTCTCAACAACAAAccatatgaatatcctccttagttcc |
| 1607 | ACATGCAGCACCGGCAGAA |
| 1608 | TGTCGCCATGTCGATACCATGC |
| 1613 | GTTTTTAATGGTTTGTTTTAAAAGTCAAAGAGGAATGAATtgtaggctggagctgcttc |
| 1614 | GAAAGAAAGGTAGTCTTCTTGACTACCTTTCGACTCTAGGccatatgaatatcctccttagttcc |
| 1615 | ACATAATGAGGTCTGAGAACG |
| 1616 | AGCCAGTTACCTTGGTTAAG |
| 1825 | ACACTCTTTCCCTACACGACGCTCTTCCGATCTCCTACGGGNGGCWGCAG |
| 1826 | GTGACTGGAGTTCAGACGTGTGCTCTTCCGATCTGACTACHVGGGTATCTAATCC |

| | |
|---|---|
| 1841 | TAATTTTAATTGATTGTTTTTAAAGTCAAAGAGGTTTTCTtgtaggctggagctgcttc |
| 1842 | AATATTTTAAAGGGAGGCAGTAACACTGCCTTCCTTTTAAccatatgaatatcctccttagttcc |
| 1843 | CCAAGTGTCTCATATCATAGT |
| 1844 | TTAACCATGACATATGCGAT |
| 1845 | GGCGAACTCTGTTATCTTGTTAACTTATAAGGAGTTATGTtgtaggctggagctgcttc |
| 1846 | ATAAAGGCTGCATAAAAAGGCCGGAACCCCGGCCCTTATAccatatgaatatcctccttagttcc |
| 1847 | TAGTTCAGGAGCTGCCCTAC |
| 1848 | ATGGCATGTACCATTGTGAGG |
| 1856 | ATTCggtaccGGCAATGAAAAGTCTTTTG |
| 1857 | TGTGctcgagCTATTTCTTGATTCCTAAACG |
| 1879 | ATCAATGCACGGTTGGCGTC |
| 1880 | GAGGAACCCTATGGTTTACTC |
| 1918 | TCCCTACGACTGCAGCTGA |
| 1919 | AATGCGGGAGGTCGAGTCCT |
| 1920 | TTCAGCACCACATGTGCCTG |
| 1921 | CAATGTGAGAGGGTTGCTG |
| 1922 | TGTGTGGCCGTCAACAGGA |
| 1923 | AGTGCGATACCTTGTGGGC |
| 2105 | ACTAAATTAATAATAATGAAAAGCTTGGAGAGTTATACCAccatatgaatatcctccttagttcc |
| 2106 | TGAGAAAAGAGCACACTGACA |
| 2107 | CATCTGACAGTCCTACCTGC |
| 2127 | TGATATATAATCTATATATAAAATAAAAAAAGGTATATATccatatgaatatcctccttagttcc |
| 2128 | TTTTTTATATCGAATGAACGACAGAAGTTGTGGAGATTTTccatatgaatatcctccttagttcc |
| 2129 | CGATGCGGTCACAATGCT |
| 2130 | GAGCATTGCTCATCGCCA |

| | |
|---|---|
| 2142 | ACACTCTTTCCCTACACGACGCTCTTCCGATCTGACACCTACGGYKCYGACAACT |
| 2143 | GTGACTGGAGTTCAGACGTGTGCTCTTCCGATCTGTCGAACTGGTACTGAGCAAC |
| 2508 | AATTCGTGTCGCTCAAGG |
| 2509 | AACAGCCAAGCTTGCATG |
| 2538 | ATTCggatccGGCAATGAAAAGTCTTTTG |
| 2771 | TGTGaagcttCTATTTCTTGATTCCTAAACG |
